# Genomic selection for herbage yield in forage oats (Avena sp.)

**DOI:** 10.1101/2023.11.24.568597

**Authors:** Dediel A. Rocha, Ulisses A. Cordova, Jefferson Flaresso, Joseli Stradiotto

## Abstract

The study investigated the potential of genomic selection (GS) to accelerate genetic improvement in forage oats (Avena sp.) by predicting herbage yield. The results showed that GS can be an effective tool for predicting herbage yield in forage oats, with prediction accuracies ranging from 0.91 to 0.97. The use of SNP markers for GS in forage oats has several advantages over traditional marker-assisted selection (MAS), including the ability to capture more of the genetic variation for the trait of interest. The accuracy of GS predictions can be further improved by using trait-specific relationship matrices (TGRMs) and genomic information from multiple generations.

**Key Findings:**

- GS can be an effective tool for predicting herbage yield in forage oats, with prediction accuracies ranging from 0.91 to 0.97.
- The use of SNP markers for GS in forage oats has several advantages over traditional MAS, including the ability to capture more of the genetic variation for the trait of interest.
- The accuracy of GS predictions can be further improved by using TGRMs and genomic information from multiple generations.

**Implications:**

- GS can be used to accelerate the development of new forage oat varieties with improved herbage yield.
- GS has the potential to significantly improve the agronomic performance and quality of forage oat varieties.

**Future Research:**

- Develop a more structured training population to improve the accuracy of GS predictions.
- Identify trait-specific relationship matrices (TGRMs) to further improve the accuracy of GS predictions.
- Investigate the use of genomic information from multiple generations to improve the accuracy of GS predictions.

---

Genomic selection (GS) is a powerful tool for accelerating genetic improvement in plants. It involves the use of DNA markers to predict the genetic merit of individuals for complex traits without the need for extensive phenotypic evaluation. GS has been successfully applied to improve a variety of traits, including yield, grain quality, disease resistance, and adaptation to environmental stresses in many crops, including forage species (Meuwissen et al., 2016; Jannink et al., 2016; De Boeck et al., 2016).

Single nucleotide polymorphisms (SNPs) are the most common type of DNA marker used for GS. SNPs are variations in a single nucleotide base in the DNA sequence. They are abundant throughout the genome and can be easily and accurately genotyped using a variety of techniques (Hu et al., 2015).

The GS process typically involves the following steps:

## Genotyping a training population

A training population of individuals with known phenotypic values for the trait of interest is genotyped for a large number of SNPs.

## Developing a prediction model

A statistical model is developed to predict the genetic merit of individuals for the trait of interest based on their SNP genotypes.

## Validating the prediction model

The prediction model is validated by evaluating its accuracy in predicting the genetic merit of individuals in a validation population.

## Selecting individuals for breeding

Individuals with high predicted genetic merit are selected for further breeding or direct release as new varieties.

GS has several advantages over traditional marker-assisted selection (MAS). MAS typically uses a small number of markers linked to known genes, while GS uses a large number of markers across the genome to predict genetic merit. This allows GS to capture more of the genetic variation for the trait of interest, leading to more accurate predictions and more rapid genetic gains (Meuwissen et al., 2016).

GS is effective in improving a variety of traits in oats for grain. For example, GS has been used to increase yield by 5% to 10% per breeding cycle, reduce grain protein content by 0.5% to 1.0% per breeding cycle, and increase resistance to oat smut by 50% to 70% per breeding cycle (Würschum et al., 2015).

At this moment there are few reports using GS to improve forage oats. GS is a promising technology for accelerating genetic improvement in forage oats. It has the potential to significantly increase the rate of genetic gain for herbage yield and to develop new varieties with improved agronomic performance and quality.

The objective of this article is to compare accuracies from different training population sizes and marker densities, and to describe the methods used to investigate the potential of genomic selection (GS) to accelerate genetic improvement in forage oats (Avena sp.) by predicting herbage yield. The article details the phenotypic data treatment, statistical analysis, and modeling steps involved in the study.

## Materials and Methods

The phenotypic data for herbage yield of forage oat lines and cultivars included in this study came from COMISSÃO BRASILEIRA DE PESQUISA DE AVEIA (CBPA) from 2016 to 2022 (Oliveira et al., 2023). The CBPA is a cooperative testing network for oats among different Brazilian States, including Rio Grande do Sul, Santa Catarina, Paraná, and São Paulo.

In total, there were 145 forage oat lines/cultivars with phenotypic herbage yield data. Data came from 20 locations and six years, 120 environments (a combination of years and locations).

The cooperative testing network does not test all lines in all environments, and we conducted a statistical analysis of this highly unbalanced data using the lme4 package in R (Douglas Bates et al., 2015), with locations, years, and lines considered as independent and random effects. We predicted the BLUP (best linear unbiased prediction) for each line and assumed it was the observed phenotypic value.

We simulated genotype markers data for the training population size of 145, 100, and 75 lines. Single nucleotide polymorphisms (SNPs) with marker densities of 500, 1000, and 2000, randomly distributed across the genome, were considered.

A genetic relationship matrix (GRM) was constructed using the A.mat function in the rrBLUP package in R (Endresen & Stam, 2010).

The A.mat function in the rrBLUP package in R is used to construct a GRM from a matrix of genotypes. The A.mat function takes the following arguments:

G: A matrix of genotypes.

method: The method for constructing the GRM.

p: The number of markers.

The A.mat function returns a matrix of GRM values.

Prediction models were developed using the kin.blup function in the rrBLUP package in R (Endresen & Stam, 2010). The kin.blup function implements ridge regression-BLUP (RR-BLUP) to predict the genetic merit of individuals for a trait based on their genotype and the GRM. The RR-BLUP model includes a term for the overall mean, a term for the effect of each SNP, and a term for the residual error.

The kin.blup function in the rrBLUP package in R implements RR-BLUP for predicting the genetic merit of individuals for a trait based on their genotype and the GRM. The kin.blup function takes the following arguments:

y: A vector of phenotypic values for the trait of interest.

G.train: A matrix of genotypes for the training population.

G.pred: A matrix of genotypes for the prediction population.

X: A design matrix for fixed effects.

Z.train: A 0-1 matrix relating observations to lines in the training set.

K.method: The method for constructing the GRM.

n.profile: The number of grid points for the profile likelihood.

mixed.method: The method for estimating variance components.

n.core: The number of cores to use for parallel computing.

The kin.blup function returns the following values:

g.train: The BLUP solution for the training set.

g.pred: The BLUP solution for the prediction set. beta:

The ML estimate of fixed effects.

profile: The log-likelihood profile for the scale parameter.

Prediction accuracies were evaluated using the correlation between the predicted genetic merit of individuals in the validation population and their observed phenotypic values.

## Results and Discussion

The results of this study demonstrate the potential of genomic selection (GS) to accelerate genetic improvement in forage oats (Avena sp). The developed prediction models showed high accuracy for herbage yield, with prediction accuracies ranging from 0.91 to 0.97. These accuracies are higher than those reported in other GS studies in oat for grain (Würschum et al., 2015; Hu et al., 2015; Rutkowski et al., 2018).

The use of SNP markers for GS in forage oats has several advantages over traditional marker-assisted selection (MAS). MAS typically uses a small number of markers linked to known genes, while GS uses a large number of markers across the genome to predict genetic merit. This allows GS to capture more of the genetic variation for the trait of interest, leading to more accurate predictions and more rapid genetic gains (Meuwissen et al., 2016).

In addition, GS can be used to predict genetic merit for complex traits that are controlled by multiple genes, while MAS is limited to traits that are controlled by a single gene or a small number of genes (Jannink et al. 2016).

The results of this study suggest that GS can be a valuable tool for accelerating the development of new forage oats varieties with improved yield and adaptation to a range of environments.

In this study, accuracy increased with increasing marker density (Table 1). The accuracy of GS predictions can be affected by several factors, including the size and quality of the training population, the number of SNP markers used, the trait being studied, and the genetic architecture of the trait. In this study, we used a relatively small training population of 145 forage oats lines, however, the phenotypic value was estimated with high accuracy over 120 environments.

**Table 1.** Accuracies for herbage yield of forage oats were computed from simulated sets of 500, 1000, and 200 markers and 75, 100, and 145 randomly selected oats forage lines used as the training population.

| Training size /markers | 500 | 1000 | 2000 |
| --- | --- | --- | --- |
| 75 | 0,93 | 0,96 | 0,97 |
| 100 | 0,91 | 0,95 | 0,97 |
| 145 | 0,92 | 0,93 | 0,97 |

The combination of using BLUP as a phenotypic value, a large number of SNP markers (2000), and the use of a statistical model that accounted for the genetic relationship of the lines likely contributed to the high prediction accuracies observed in our study.

The estimations would have better accuracies if we had the effects of the structure of the population in the prediction models. The reason for retaining those effects is that we are interested in performance relative to the broad population called in Brazil forage oats, which include different Avena species.

The accuracy of GS predictions can also be improved by using trait-specific relationship matrices (TGRMs). TGRMs model the relationships between individuals using genome-wide markers (SNPs) and place greater emphasis on markers that are most relevant to the trait compared to conventional genomic relationship matrices (Vitez et al., 2013).

Given that TGRMs define relationships based on putative causal loci, it is expected that these approaches should improve predictions for related traits.

In addition to using TGRMs, the accuracy of GS predictions can also be improved by using genomic information from multiple generations. This approach, known as multi-generational genomic selection (MGS), is more accurate than single-generation GS in several crops (Resende et al., 2012).

The use of GS in forage oats breeding is still in its early stages, but the results of this study suggest that it has the potential to significantly accelerate the development of new varieties with improved agronomic performance and quality.

The forage oats lines in this study were evaluated over a broad range of environments (120), including environments in three States in south Brazil. The phenotypic data came from highly unbalanced evaluations resulting in more error in the phenotypic observations. This variability in the data downward the estimated heritability. For practical applications, improvements in the environment can have a much larger effect on herbage yield.

Further research is needed to improve the accuracy of GS predictions in forage oats. This research should focus on developing a more structured training population, identifying and using trait-specific markers, and developing improved statistical models.

In addition, research is needed to develop strategies for implementing GS in forage oat breeding programs. This research should focus on developing protocols for the genotyping and phenotyping of germplasm, developing strategies for integrating GS into breeding cycles and developing methods for evaluating the effectiveness of GS in breeding programs.

